## Supplemental Figures and Text for "Sea anemone genomes reveal ancestral metazoan chromosomal macrosynteny"

### Supplementary information for *Sea anemone genomes reveal ancestral metazoan chromosomal macrosynteny*


1. **Supplementary Figures**
    1.1. Divergence time estimates of cnidarians
    1.2. Genome size, assembly quality and assembly completeness estimates
    1.3. Alignments of candidate *N. vectensis* pseudo-chromosome assemblies
    1.4. Summary of *N. vectensis* gene models
    1.5. Locations of ANTP class homoebox complements on the *E. muelleri* and *N. vectensis* genome
    1.6. Contact maps of two largest chromosomes of the *N. vectensis* assembly
    1.7. Cnidarian ancestral linkage groups
    1.8. Bilaterian ancestral linkage groups
    1.9. Metazoan ancestral linkage groups
    1.10. NJ and ML analysis of the NK cluster and SuperHox cluster proteins
    1.11. ML analysis of the NK-like proteins including the NK-like proteins from the earlier branching clades
    1.12. Non-bilaterians have less pronounced TAD boundaries than bilaterians
    1.13. Summary of orthologous groups used in macrosynteny detection and ALG inference
    1.14. Inference of ancestral linkage groups
2. **Supplementary Text**
    2.1. Assembly of *Nematostella vectensis* and *Scolanthus callimorphus* genomes
    2.2. Chromosomal repeat dynamics in the Edwardsiidae
    2.3. Evolution the homeobox gene clusters in early branching metazoans
    2.4. Ultraconserved non-coding Elements
    2.5. The SIMRBase Genome Browser
    2.6. Inference of ancestral linkage groups
3. **Supplementary Tables**
    3.1. Summary of repetitive sequence content in the *N. vectensis* and *S. callimorphus* genomes reported by RepeatMasker
    3.2. Chromosome-wise abundance and enrichment of repeat classes in *N. vectensis*
    3.3. Chromosome-wise abundance and enrichment of repeat families in *N. vectensis*
    3.4. Chromosome-wise abundance and enrichment of repeat classes in *S. callimorphus*
    3.5. Chromosome-wise abundance and enrichment of repeat families in *S. callimorphus*
    3.6. Genomic locations of a subset of the ANTP, PRD, SINE, TALE and HNF class homeobox genes in the *N. vectensis* genome
    3.7. Table of correspondence between *N. vectensis* cloned complete CDS sequences in the NCBI database and NV2 models

3.8. Ultraconserved Noncoding Elements in the *N. vectensis* genome
3.9. Genes contained in the cnidarian ancestral linkage groups
3.10. Genes contained in the bilaterian ancestral linkage groups
3.11. Genes contained in the metazoan ancestral linkage groups
3.12. Number of microsyntenic blocks in deuterostomes, spiralians and cnidarians

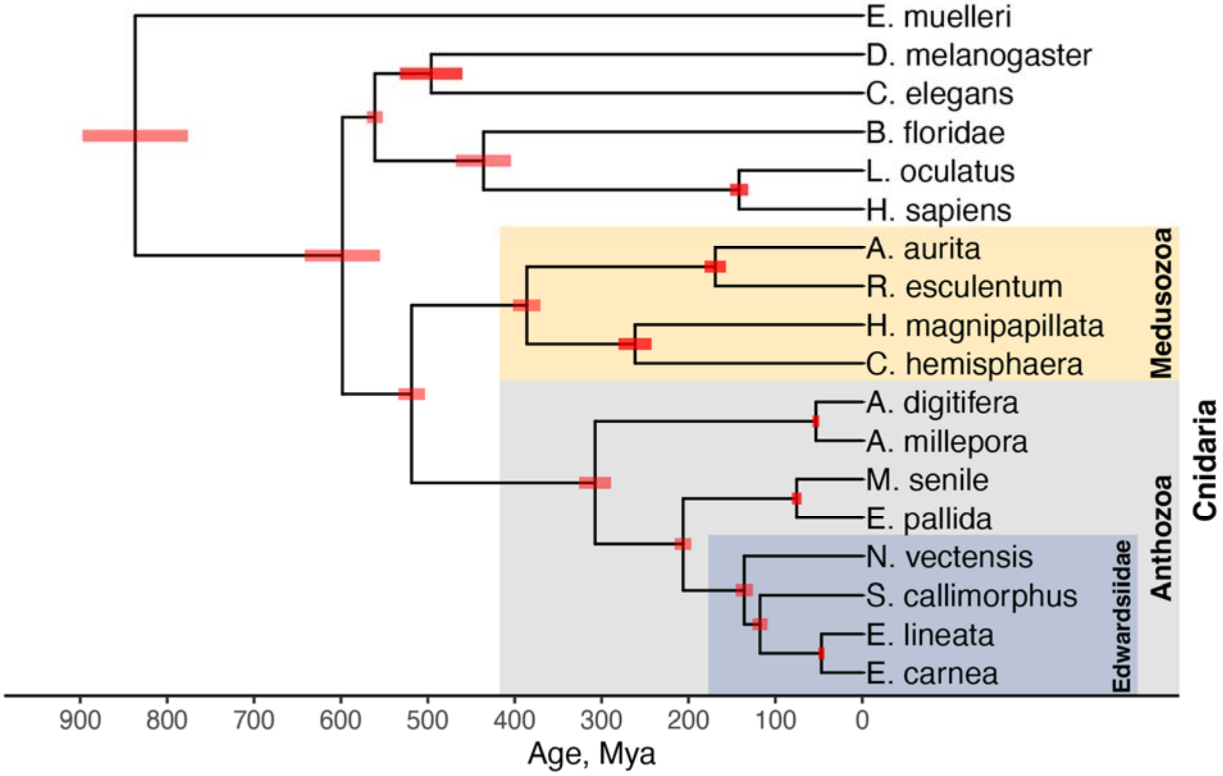


**Extended Data Figure 1.** Divergence time estimates of cnidarians, calibrated by an estimate of the cnidarian-bilaterian split between 595.7 and 688.3 Mya[1].

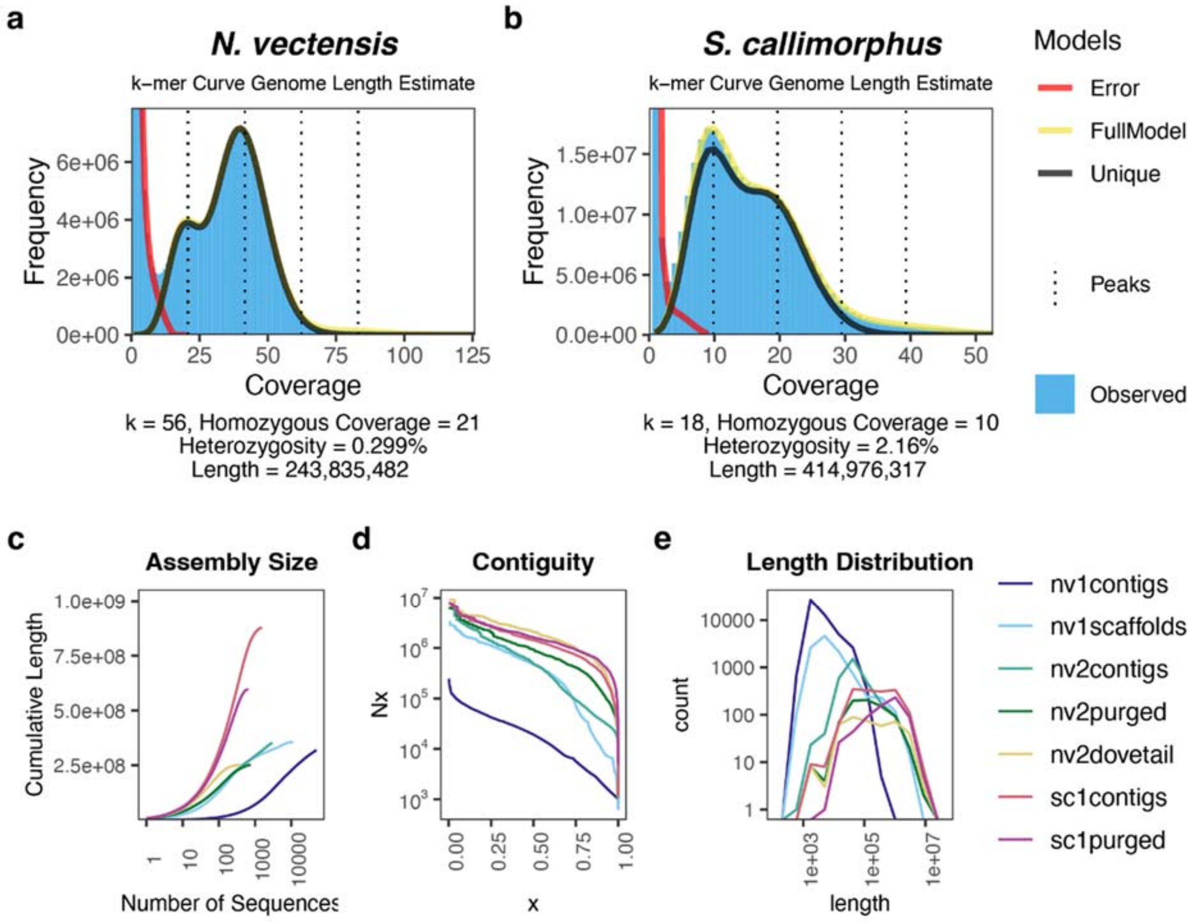


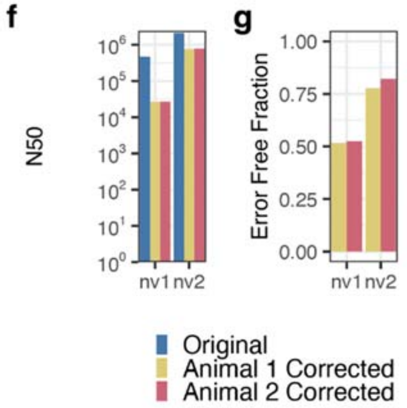


h

| Assembly | Level | Length | No. of Sequences | Median Length | N50 |
|---|---|---|---|---|---|
| nv1 | contigs | 318,900,652 | 48,580 | 2,373 | 19,187 |
| | scaffolds | 356,613,585 | 10,804 | 6,708 | 472,588 |
| nv2 | contigs | 351,276,352 | 2,951 | 38,162 | 488,093 |
| | purged | 252,158,603 | 751 | 107,017 | 938,720 |
| | dovetail | 252,196,803 | 409 | 129,750 | 2,101,135 |
| | chromosomes | 247,748,268 | 15 | 16,174,719 | |
| sc1 | contigs | 879,055,531 | 1,517 | 212,192 | 1,435,147 |
| | purged | 596,774,616 | 645 | 623,295 | 1,613,650 |
| | chromosomes | 452,358,050 | 15 | 28,798,654 | |

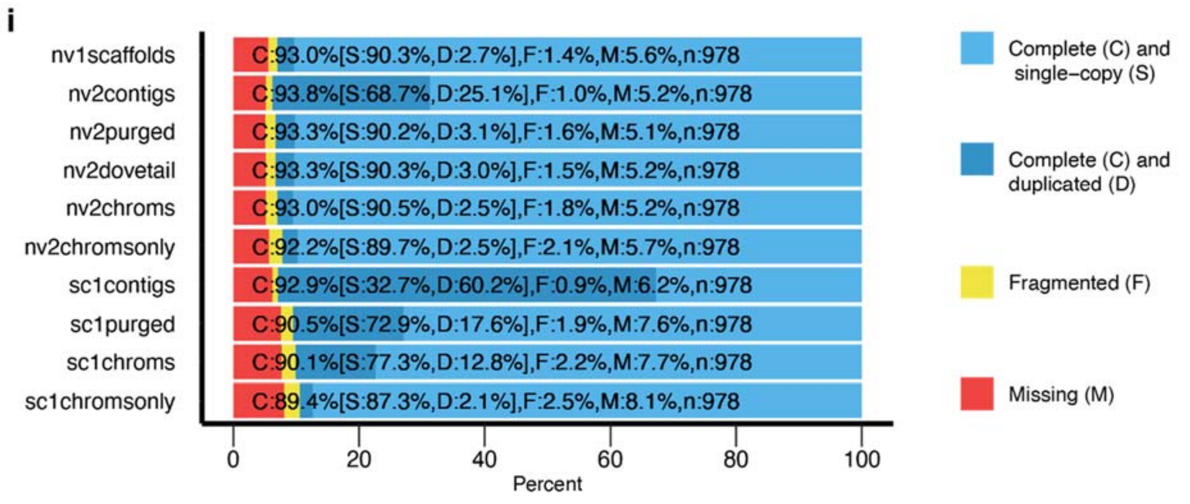

**Extended Data Figure 2.** Quality assessment of the previous *Nematostella vectensis* assembly (nv1)[2], the assembly presented in this paper (nv2) and *Scolanthus callimorphus* (sc1) genomes at different assembly levels. Contigs refer to sequences which have been contiguously assembled by read overlap. “Purged” assemblies have been filtered against redundant contigs appearing due to heterozygosity (see Materials and Methods). Scaffolds and “dovetail” are collections of contigs which have been ordered by evidence from BAC ends (scaffolds) or in vitro proximity ligation (“dovetail”) and “chromosomes” are collections of contigs (sc) or dovetail scaffolds (nv) ordered by in vivo proximity ligation. a,b) *k*-mer curve genome size and heterozygosity estimates of short read data from *N. vectensis* and *S. callimorphus*. c) Assembly size characterized by the cumulative length as a function of number of sequences d) Nx contiguity c) sequence unit (contig, scaffold or dovetail scaffold) length distribution. f,g) Assembly fidelity of the nv1 and nv2 assemblies with respect to intra-scaffold and intra-contig assembly correctness, assessed by the sequenced paired-end reads of 2 individuals and analyzed using the REAPR pipeline. f) *N. vectensis* assembly N50 after stringent breaking of dovetail scaffolds. A smaller reduction from the “original” N50 indicates a better assembly. g) The estimated error-free fraction, in terms of sequence fidelity and contiguity, of the original nv1 and nv2 genomes. h) Summary statistics of the nv1, nv2 and sc1 genomes at all levels. Lengths at the “chromosome” level indicate the total length of the 15 pseudo-chromosomes including a 100 base gap at each of the scaffold junctions. i) Estimates of assembly completeness and redundancy by proxy of conserved pan-metazoan single-copy genes (“BUSCOs”). “chromsonly” assemblies exclude unplaced dovetail scaffolds (nv) or contigs (sc) and only include the respective 15 chromosomes.

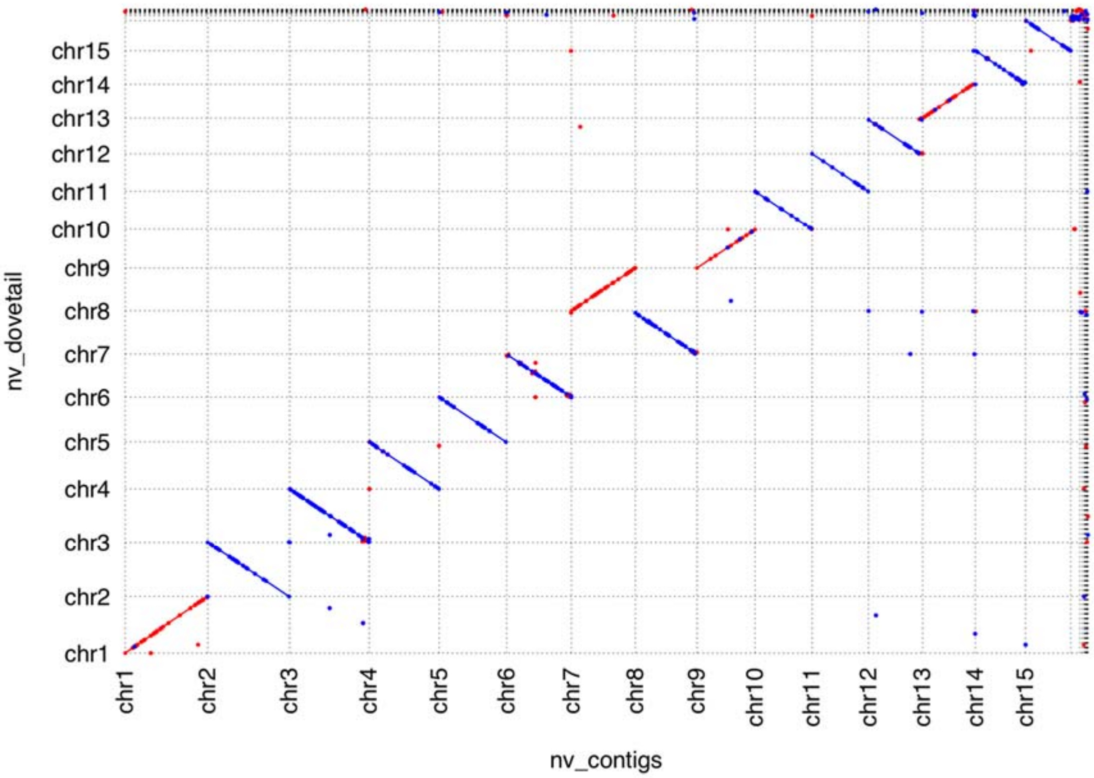


**Extended Data Figure 3.** Genome-genome alignment of two *N. vectensis* chromosome level assemblies. Red represents forward alignments and blue represents reverse alignments. The assembly on the *x*-axis was generated using the contigs after purging haplotigs as a starting point, and the assembly on the *y*-axis using the Dovetail-scaffolded contigs as a starting point. Chromosomes are labeled and unplaced contigs or scaffolds are placed after chr15. Both were independently and blindly subjected to manual review.

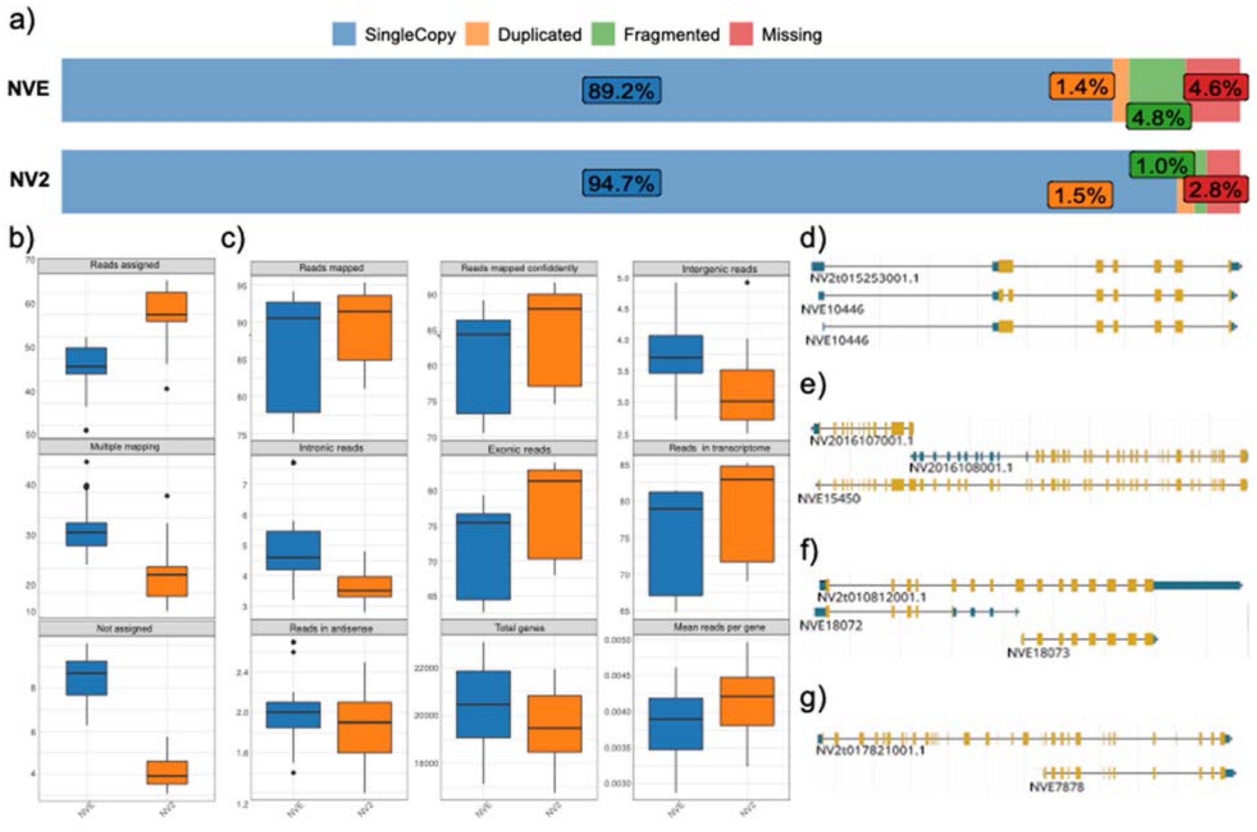


**Extended Data Figure 4. Comparison of NVE and NV2 gene models;** Comparison of NVE and NV2 gene models. a) Busco completeness using the Metazoan ODB10 database shows an increase of 5% in full single copy orthologs and a reduction of missing and fragmented orthologs in the NV2 annotation; b) Percentage of bulk RNA-seq reads unequivocally assigned to a single gene model (Assigned), to multiple gene models (Multiple mapping) and to intergenic regions (Not assigned) in NVE and NV2 models; c) Comparison of single cell RNAseq alignments to the NVE and NV2 genomes. On average 10% more reads can be unequivocally assigned to NV2 genes models compared to NVE models. The total number of genes identified in the NVE genome is greater than in the NV2 genome. This is due to the presence of multiple uncollapsed isoforms in the NVE genome and more fragmented gene models as seen in the BUSCO results and in the average number of reads per gene plot; d) example of uncollapsed NVE model that are actually one single NV2 model in chr3; e) example of 1 NVE model split into 2 NV2 models based on Isoseq data on chr4; f) example of two non-overlapping NVE models merged into a single NV2 model on chr2; g) example of a model with an extended 5'-end on chr5

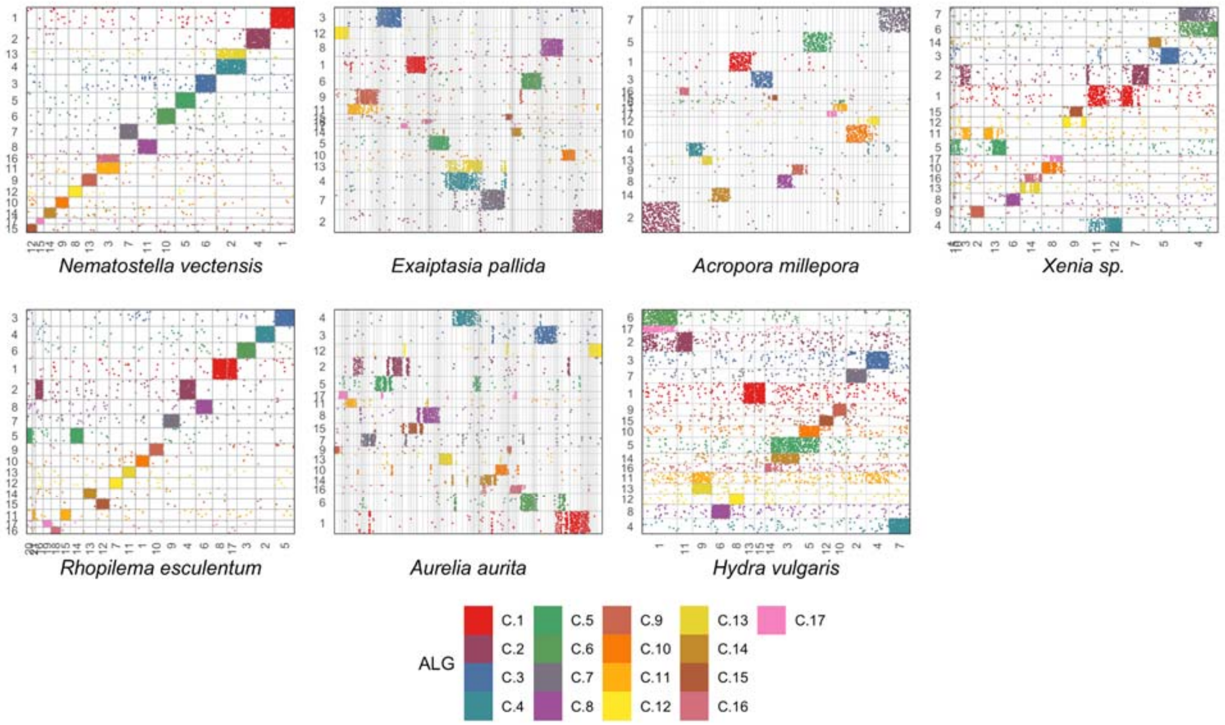


**Extended data figure 5.** Dot plots representing positions of genes from ancestral cnidarian linkage groups (ALGs) in extant chromosomes. The *y*-axis represents the 19 ALGs from the cnidarian lineage and dots are redundantly colored for clarity. The *x*-axis represents the extant genome. In the case of the genomes assembled only at the scaffold level, no chromosomes are represented on the *x*-axis. Scaffold order in the dot plots is determined using hierarchical clustering of the ALG-wise scaled matrix of genes shared between the extant genome and the ALG. A complete list of genes can be found in EDT 9.

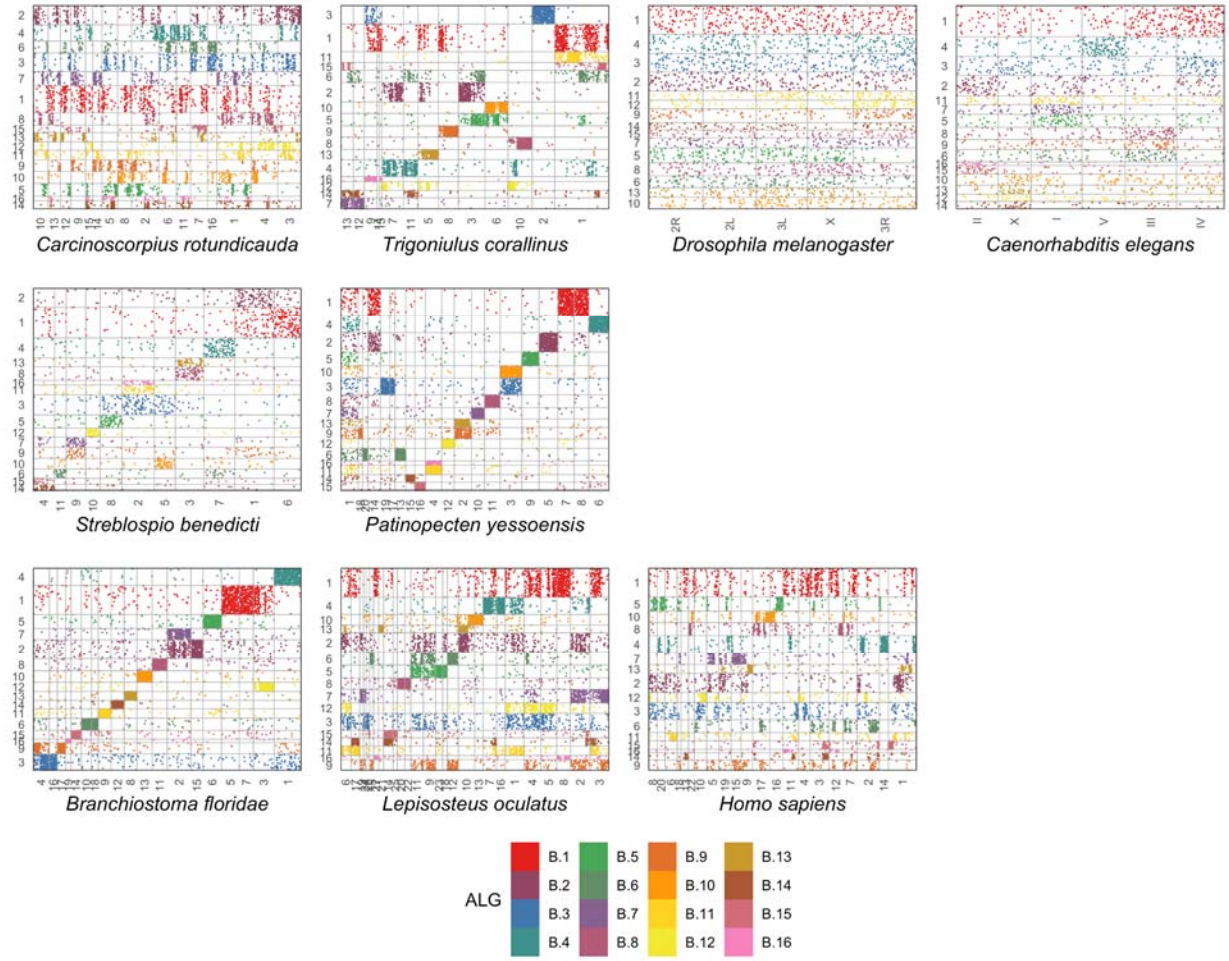


**Extended data figure 6.** Same analysis as EDF 5 using the bilaterian ancestral linkage groups. A complete list of genes can be found in EDT 10.

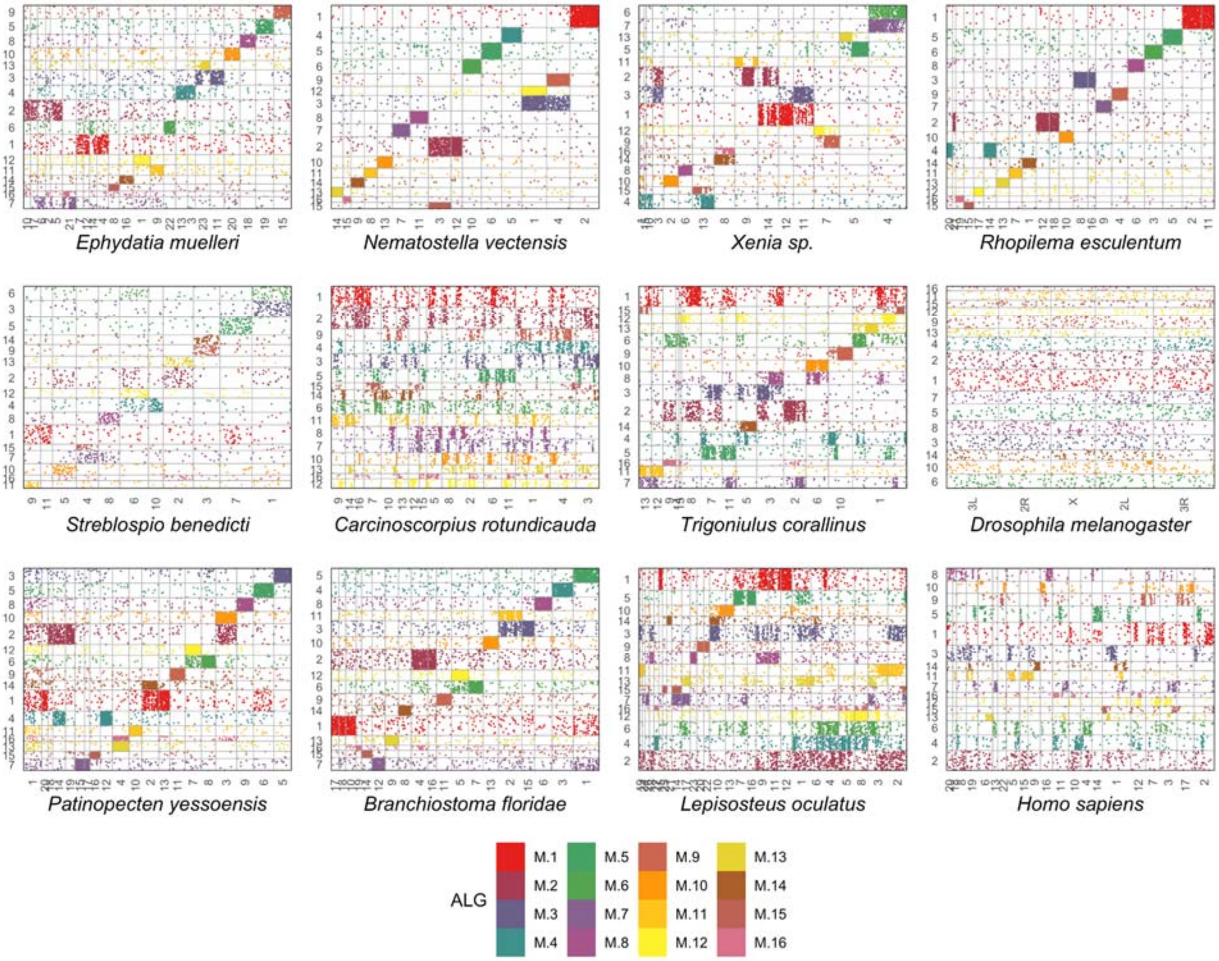


**Extended data figure 7.** Same analysis as EDF 5 using the metazoan ancestral linkage groups. A complete list of genes can be found in EDT 11.

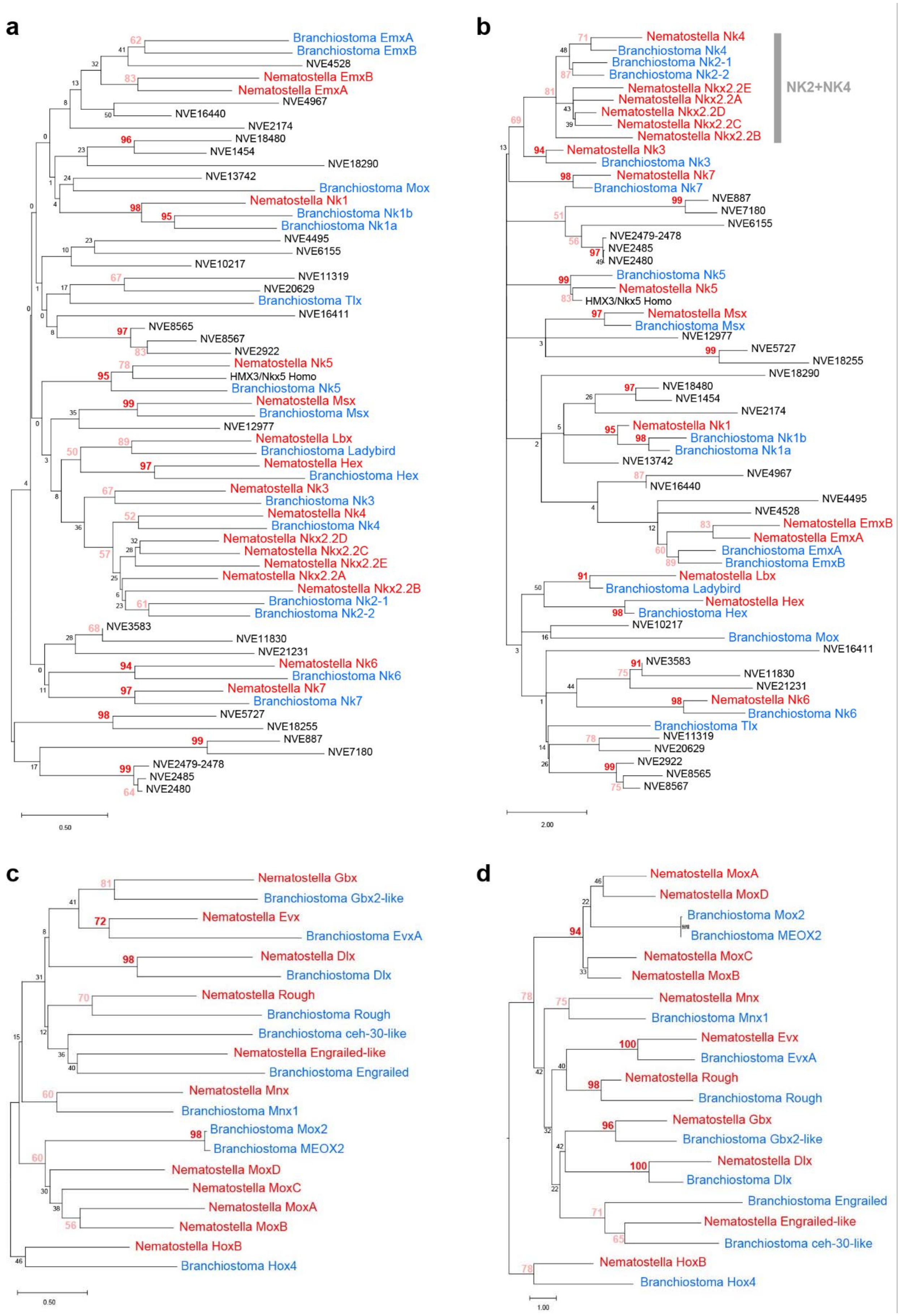
a
Branchiostoma EmxA
Branchiostoma EmxB
NVE4528
Nematostella EmxB
Nematostella EmxA
NVE4967
NVE16440
NVE2174
NVE18480
NVE1454
NVE18290
NVE13742
Branchiostoma Mox
Nematostella Nk1
Branchiostoma Nk1b
Branchiostoma Nk1a
NVE4495
NVE6155
NVE10217
NVE11319
NVE20629
Branchiostoma Tlx
NVE16411
NVE8565
NVE8567
NVE2922
Nematostella Nk5
HMX3/Nkx5 Homo
Branchiostoma Nk5
Nematostella Msx
Branchiostoma Msx
NVE12977
Nematostella Lbx
Branchiostoma Ladybird
Nematostella Hex
Branchiostoma Hex
Nematostella Nk3
Branchiostoma Nk3
Nematostella Nk4
Branchiostoma Nk4
Nematostella Nkx2.2D
Nematostella Nkx2.2C
Nematostella Nkx2.2E
Nematostella Nkx2.2A
Nematostella Nkx2.2B
Branchiostoma Nk2-1
Branchiostoma Nk2-2
NVE3583
NVE11830
NVE21231
Nematostella Nk6
Branchiostoma Nk6
Nematostella Nk7
Branchiostoma Nk7
NVE5727
NVE18255
NVE887
NVE7180
NVE2479-2478
NVE2485
NVE2480
0.50
b
Nematostella Nk4
Branchiostoma Nk4
Branchiostoma Nk2-1
Branchiostoma Nk2-2
Nematostella Nkx2.2E
Nematostella Nkx2.2A
Nematostella Nkx2.2D
Nematostella Nkx2.2C
Nematostella Nkx2.2B
NK2+NK4
Nematostella Nk3
Branchiostoma Nk3
Nematostella Nk7
Branchiostoma Nk7
NVE887
NVE7180
NVE6155
NVE2479-2478
NVE2485
NVE2480
Branchiostoma Nk5
Nematostella Nk5
HMX3/Nkx5 Homo
Nematostella Msx
Branchiostoma Msx
NVE12977
NVE5727
NVE18255
NVE18290
NVE18480
NVE1454
NVE2174
Nematostella Nk1
Branchiostoma Nk1b
Branchiostoma Nk1a
NVE13742
NVE4967
NVE16440
NVE4495
NVE4528
Nematostella EmxB
Nematostella EmxA
Branchiostoma EmxA
Branchiostoma EmxB
Nematostella Lbx
Branchiostoma Ladybird
Nematostella Hex
Branchiostoma Hex
NVE10217
Branchiostoma Mox
NVE16411
NVE3583
NVE11830
NVE21231
Nematostella Nk6
Branchiostoma Nk6
Branchiostoma Tlx
NVE11319
NVE20629
NVE2922
NVE8565
NVE8567
2.00
c
Nematostella Gbx
Branchiostoma Gbx2-like
Nematostella Evx
Branchiostoma EvxA
Nematostella Dlx
Branchiostoma Dlx
Nematostella Rough
Branchiostoma Rough
Branchiostoma ceh-30-like
Nematostella Engrailed-like
Branchiostoma Engrailed
Nematostella Mnx
Branchiostoma Mnx1
Branchiostoma Mox2
Branchiostoma MEOX2
Nematostella MoxD
Nematostella MoxC
Nematostella MoxA
Nematostella MoxB
Nematostella HoxB
Branchiostoma Hox4
0.50
d
Nematostella MoxA
Nematostella MoxD
Branchiostoma Mox2
Branchiostoma MEOX2
Nematostella MoxC
Nematostella MoxB
Nematostella Mnx
Branchiostoma Mnx1
Nematostella Evx
Branchiostoma EvxA
Nematostella Rough
Branchiostoma Rough
Nematostella Gbx
Branchiostoma Gbx2-like
Nematostella Dlx
Branchiostoma Dlx
Branchiostoma Engrailed
Nematostella Engrailed-like
Branchiostoma ceh-30-like
Nematostella HoxB
Branchiostoma Hox4
1.00

**Extended Data Figure 8. NJ and ML analysis to confirm the identity of the NK cluster and SuperHox cluster proteins.** (a) NJ (JTT+G4, bootstrap 100) and (b) ML (JTT+I+F+G4, bootstrap 100) analyses show that *Nematostella* possesses clear orthologs of *Nk1*, *Nk5*, *Msx*, *NK4*, *Nk3*, *Nk7*, *Nk6*, *Hex*, *Lbx*, as well as four *NK2.2* genes. Another *NK* gene, *NVE10217*, with some similarity to *Tlx* according to BLAST, is located on the same chromosome as all the other *NK* genes except the *NK2.2* group. However its orthology to bilaterian *Tlx* is not supported by either of the trees. (c-d) NJ (c) and ML (b) analyses show that *Nematostella* possesses clear orthologs of the non-Hox/ParaHox members of the SuperHox cluster *Mox* (four *Mox* paralogs are present in *Nematostella*), *Rough*, *Mnx*, *Evx*, *Gbx*, and *Dlx*, as well as a likely *Engrailed* ortholog. *Nematostella HoxB* and *Branchiostoma Hox4* were used as an outgroup to the non-Hox/ParaHox genes. Bootstrap values higher than 90% are shown in red; bootstrap values higher than 50% are shown in pink. Black "NVE" gene models (https://figshare.com/articles/Nematostella_vectensis_transcriptome_and_gene_models_v2_0/807696) are *Nematostella* gene models of NK-like genes with unclear orthology.

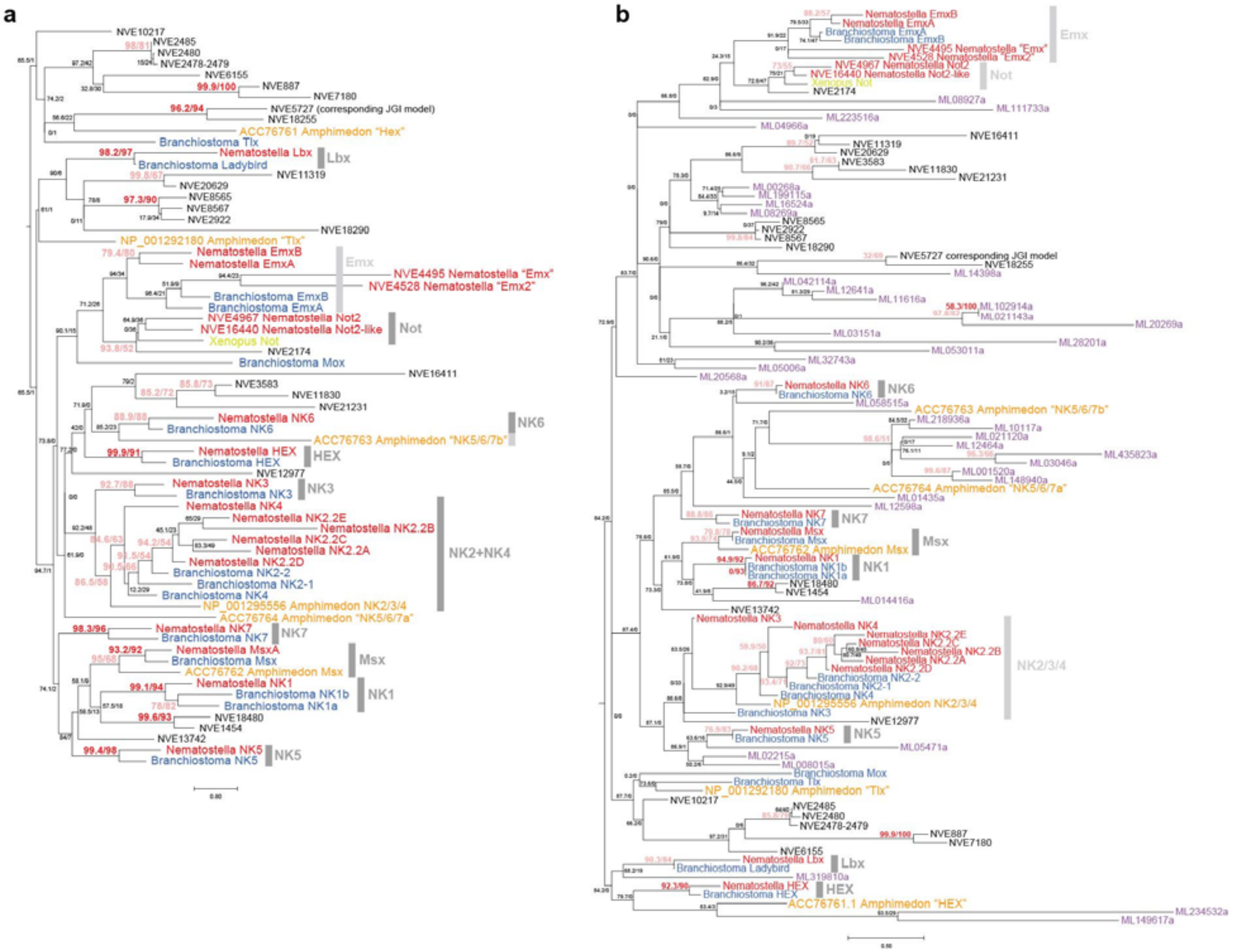


**Extended Data Figure 9. ML analysis of the NK-like proteins including the NK-like proteins from the earlier branching clades.** (a) ML tree (Q.insect+F+I+G4, bootstrap 100, aLRT 1000) containing NK-like protein sequences from the sponge *Amphimedon queenslandica* confirms orthology of the Nematostella NK cluster proteins proposed by the analyses shown on EDF 10a-b. Among *Amphimedon queenslandica* proteins, only Msx, NK2/3/4 (which groups together with the NK2/NK4 proteins), and, possibly, “NK5/6/7b” and “HEX” clearly group together with their suggested Nematostella and Branchiostoma orthologs. (b) ML tree (Q.insect+G4, bootstrap 100, aLRT 1000) containing NK-like protein sequences from the sponge *Amphimedon queenslandica* and the ctenophore *Mnemiopsis leydii* (purple “ML” protein models) shows that addition of the ctenophore sequences does not affect the grouping of the confirmed cnidarian and bilaterian NK orthologues. However, ctenophore proteins do not appear to belong to any of the NK or NK-like orthology groups from Cnidaria+Bilateria. Numbers at the nodes: bootstrap 100SH-aLRT support (%) / ultrafast bootstrap support (%).  Bootstrap values higher than 90% are shown in red; bootstrap values higher than 50% are shown in pink.

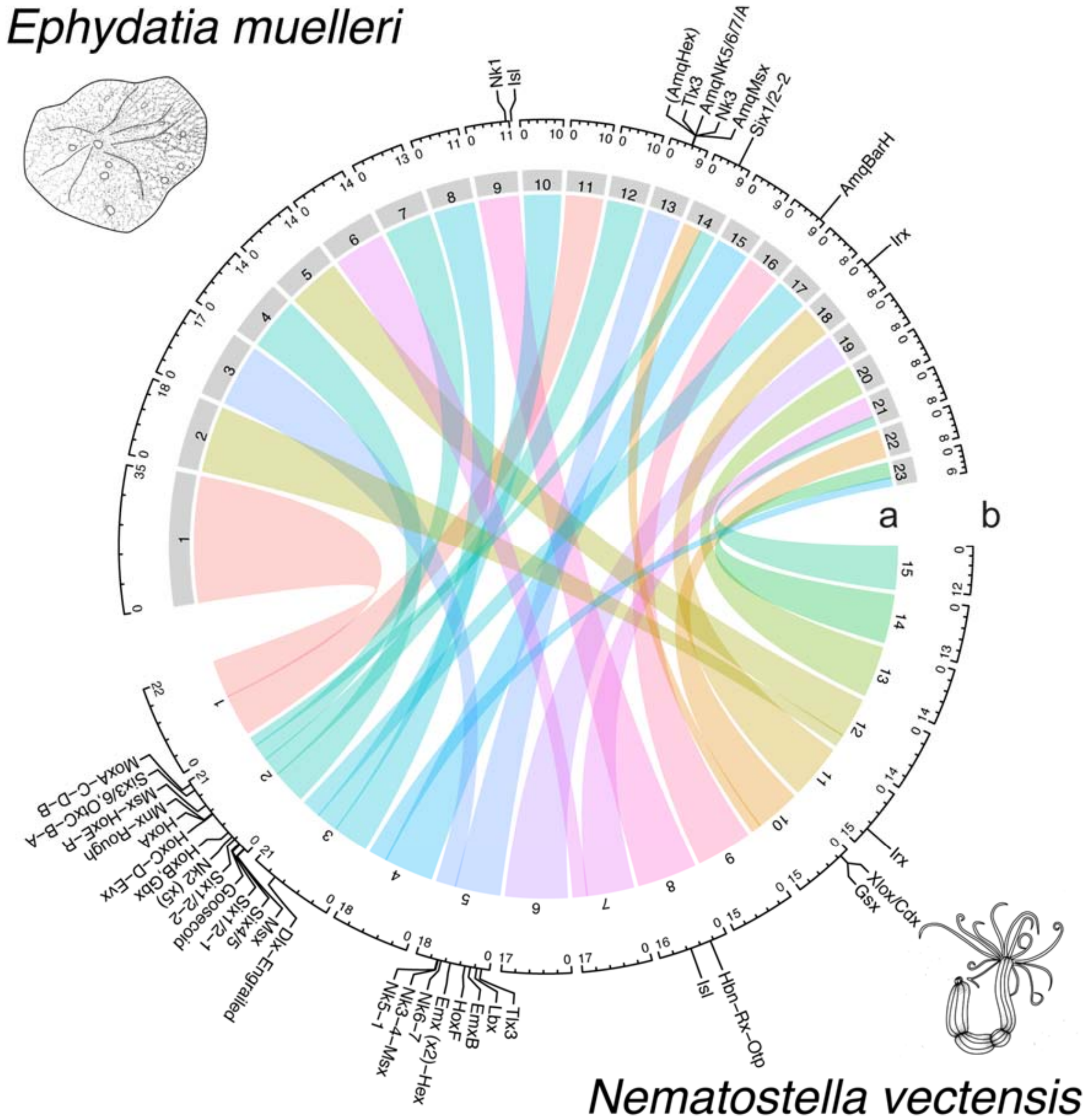


**Extended data figure 10. Locations of Hox genes and relationships of *N. vectensis*, and *Ephydatia muelleri* chromosomes.** a) Chord diagram of macrosyntenic relationships of chromosomes. b) Locations of NK and extended Hox cluster genes on the genomes.

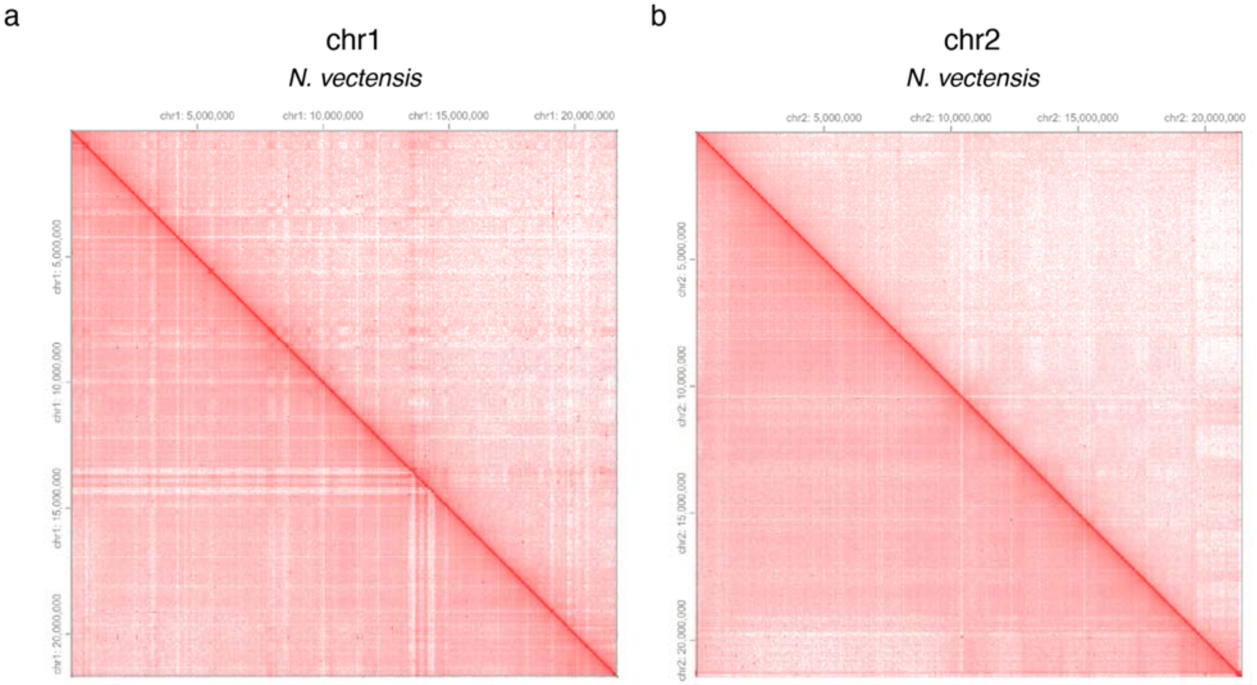


**Extended Data Figure 11. Chromosome-level Hi-C contact maps.** Contact maps for the two largest chromosomes of the *Nematostella* assembly generated by HiGlass[3] at 40k resolution. Each point represents a binned, normalized intensity of chromosomal contact as measured by the number of ligated fragments sequenced. The upper triangle represents the Hi-C experiment from this study and the lower triangle represents an external data set[4].

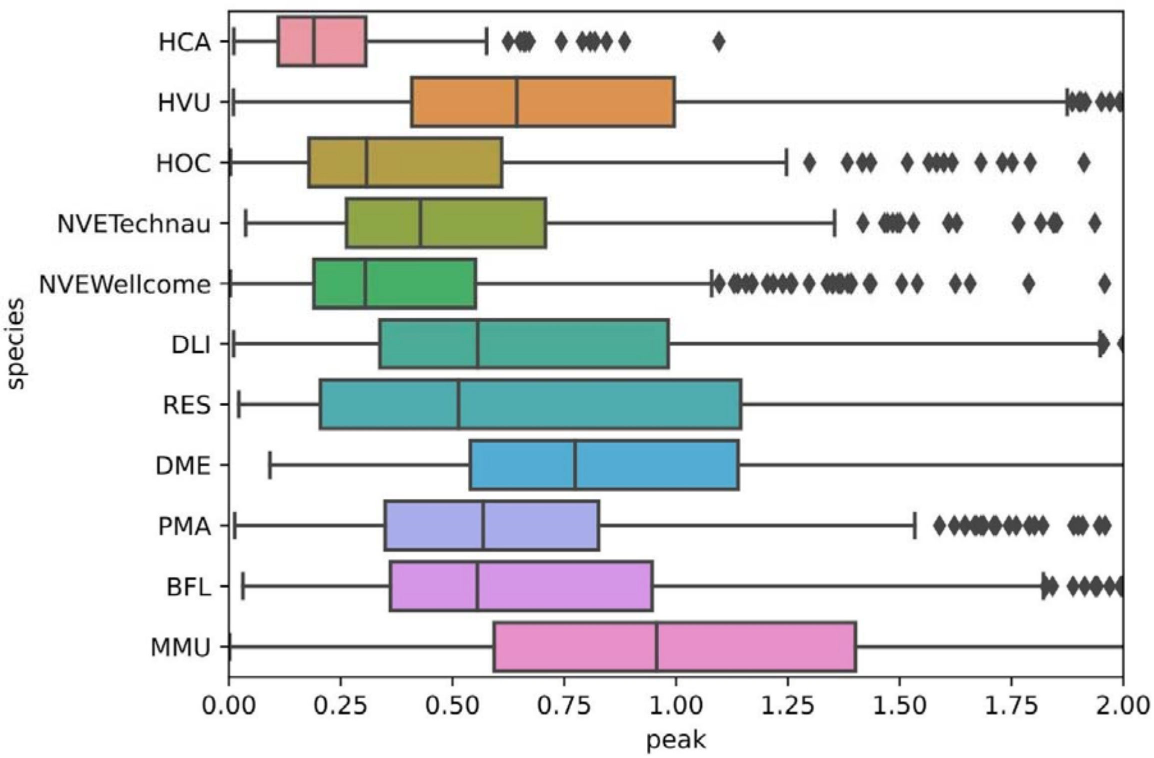


**Extended Data Figure 12**. **Non-bilaterians have less-pronounced TAD boundaries than bilaterians.** The distribution of tad peak-valley insulation score differences in animals. Ctenophores (Hormiphora californensis - HCA), Cnidarians (Hydra vulgaris - HVU, Haliclystus octoradiatus - HOC, Nematostella vectensis - NVE, Diadumene lineata - DLI, Rhopilema esculentum - RES), Bilaterians (Drosophila melanogaster - DME, Pecten maximus - PMA, Branchiostoma floridae - BFL, Mus musculus - MMU).

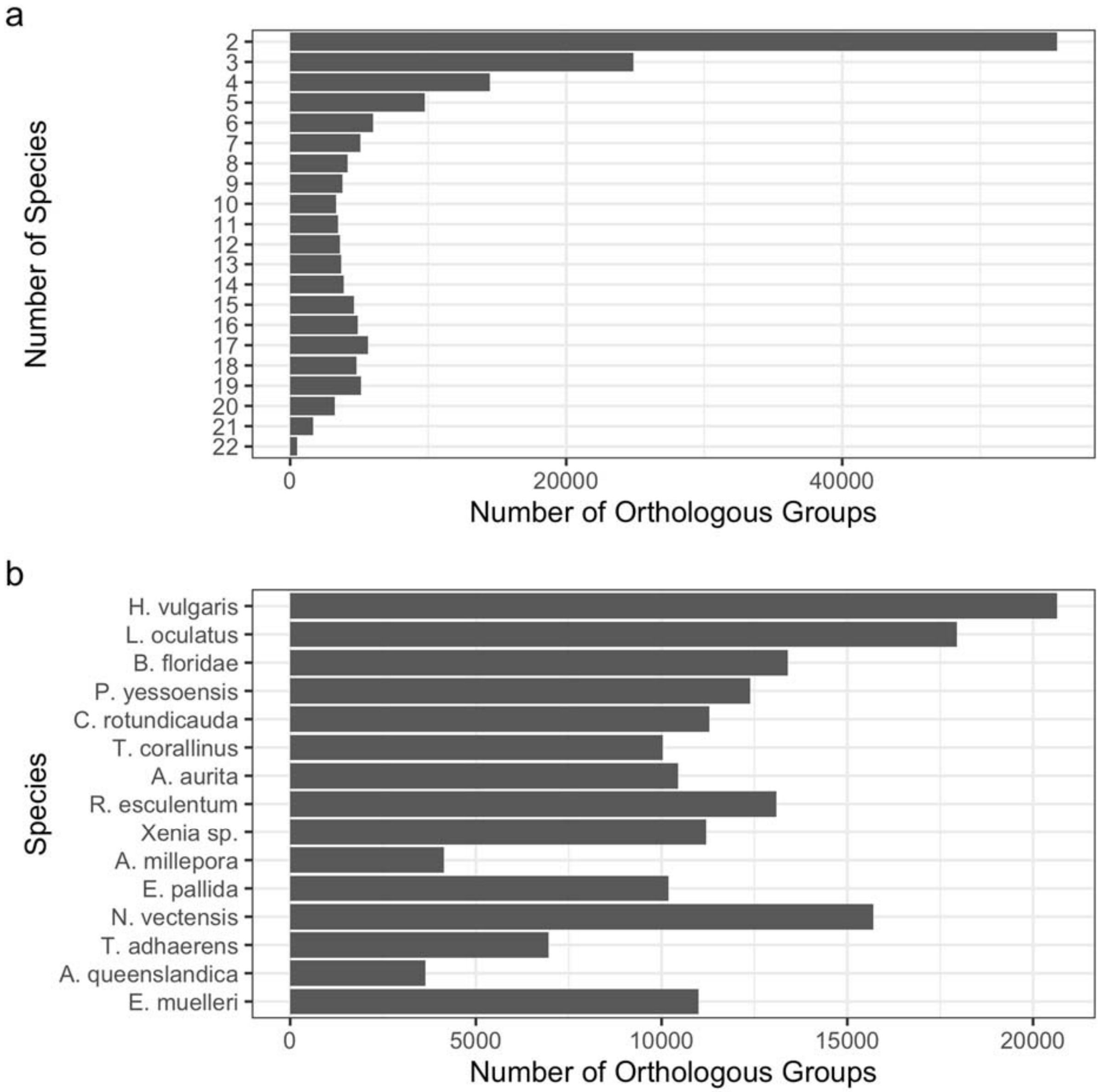


**Extended Data Figure 13. Summary of orthologous groups used in macrosynteny detection and ALG inference.** OMA orthogroups were inferred as described in the supplementary text. a) Histogram of the number of orthology groups of differing sizes. Only one gene per species was allowed, so the number of species implies the number of genes in the group. b) Total number of genes included in orthologous groups in each genome.

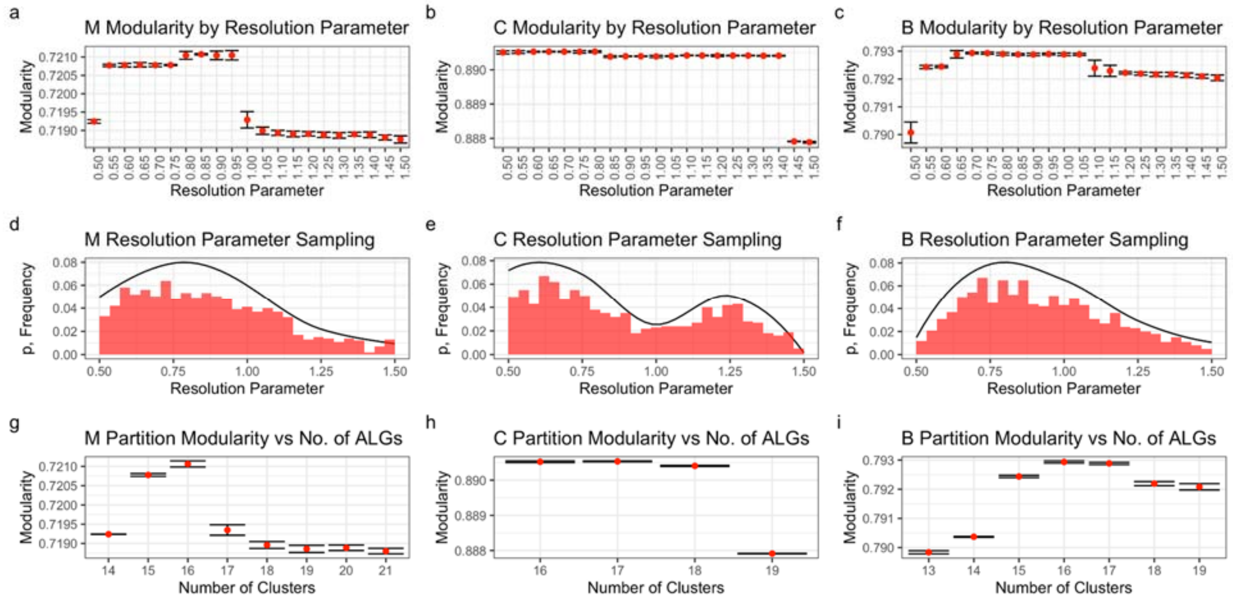


**Extended Data Figure 14. Inference of ancestral linkage groups for Metazoa (M), Cnidaria (C) and Bilateria (B).** a-c) Each setting of the resolution parameter was run with the given setting 100 times and the resulting distributions of modularity values for the partitions are represented here. The dot represents the mean modularity and the error bars +/- 1 standard deviation. d-f) A distribution of the resolution parameter was generated using the ranks of all modularity values (see supplementary text for details). The line represents the distribution from which 1000 resolution parameters were sampled. The bars are a frequency histogram of the samples taken from the distribution. g-i) 1000 partitions were generated using the Leiden algorithm using the sampled resolution parameters. Results were grouped by the number of ALGs (Clusters) inferred, and the distributions of modularity are again represented as a dot for the mean and the error bars +/- 1 standard deviation.

#### Supplementary Text

##### 1. Assembly of *Nematostella vectensis* and *Scolanthus callimorphus* genomes

*Nematostella* and *Scolanthus* PacBio long-read libraries were constructed from the same DNA used to estimate the genome size (*Scolanthus*) or individuals from the same clonal population (*Nematostella*). Self-corrected PacBio sequences were assembled into initial contigs which were already highly contiguous after an initial pass (nv2contigs, sc1contigs, EDF 2c-e). The initial assembly showed indications of redundancy as indicated by Benchmarking of Universal Single Copy Orthologs (BUSCO) scores (EDF 2i), likely caused by heterozygous alleles assembling into separate contigs. The greater number of duplicate BUSCOs in *Scolanthus* corresponds to its higher heterozygosity as indicated by the k-mer model (EDF 2a-b). After removal of these redundant contigs, the duplicate BUSCOs were reduced 3.4-fold in *Scolanthus* and 8-fold in *Nematostella* (EDF 2i).

Compared to the published *Nematostella* assembly[2], the contiguity of both genomes in terms of contig-level N50 was over 25 fold higher (EDF 2c-e,h). The *Nematostella* assembly was further scaffolded by generating libraries using the Dovetail Chicago in vitro proximity ligation platform (see Materials and Methods for details). The Dovetail-scaffolded genome further increased the N50 contiguity by 2-fold (EDF 2h). In addition, one BUSCO gene match which was previously fragmented due to an assembly break in the contig-level assembly was united on a single scaffold in the new Dovetail assembly.

We were additionally able to validate the order and correctness of the *Nematostella* intra-scaffold sequence using the REAPR pipeline[5]. We extracted DNA fragments from two individuals, measured the insert size distribution and sequenced paired ends. REAPR was used to break the genome where a substantial portion of paired read mappings on the contigs conflicted with the expected distance. The contiguity of the broken assemblies, as compared to the initial scaffolded assemblies, was much higher than that of the original *Nematostella* assembly, and also relatively higher as a fraction of the raw N50 (EDF 2f). Additionally, the fraction of the scaffolded genome considered to be error-free (in terms of both sequence and contiguity) was 150% and 157% of the previous *Nematostella* scaffolds based on the data from each of the individuals (EDF 3g). Taken together, not only were the nv2 scaffolds substantially more contiguous than the previous nv1 scaffolds[2], the sequences within these scaffolds exhibited fewer misassemblies and errors (EDF2i).

In order to obtain a chromosome-level assembly of *Nematostella* and *Scolanthus*, we performed high throughput chromosomal conformation capture (Hi-C) on a single individual from each species. In the case of *Nematostella*, the sex was male, for *Scolanthus* it was unknown. After automated assembly followed by assembly review (see Materials and Methods for details). The HiC contact maps showed evidence for 15 chromosomes for both *Nematostella* and *Scolanthus* (Figure 1d,e). This is in line with the previous estimates based on the analyses of *Nematostella* chromosome spreads[2,6]. We were also able to validate the Dovetail scaffolding by performing a second, independent assembly of the purged contigs using our Hi-C data. As shown in EDF 3, the assemblies have very long segments of identical sequence in the same arrangement and orientation, confirming the robustness of both the scaffolding and the assembly method. The minor differences in these assemblies were inspected in further manual review and ties were broken according to the Hi-C contact signal in the Dovetail assembly.

For *Nematostella*, we previously sequenced several transcriptome libraries from various developmental stages, which were assembled earlier as “NVE” gene models[7]. To further improve the gene annotation, in particular with respect to isoforms and untranslated regions, we used a combination of IsoSeq and RNAseq data, which allowed us to identify 24,525 gene models and 36,280 transcripts, termed NV2 gene models. BUSCO analysis showed that the NV2 transcriptome contains 96.1% of expected metazoan single-copy sequences, which represents an increase over the previous NVE gene models (90.6% complete BUSCOs) (EDF 4). Additional comparison to previously cloned complete CDS (EDT 7) showed that 261 of the 277 sequences (94.2%) were present. Of the missing 16 sequences, 15 could be confidently aligned to the reference genome and have been manually added to the annotation files. While the total number of NV2 gene models is similar to the NVE models, it demonstrates that many NVE models are split or fused versions of two distinct genes. Moreover, due to the long 3' UTR regions of the IsoSeq sequences, the NV2 models map in general more faithfully to single cell data (Alison). In addition, In order to compare the genomic location of homologous genes between the edwardsiids, we predicted 24,625 gene models in the *Scolanthus* genome using the previously sequenced transcriptome[8] (see Materials and Methods for details).

To showcase the quality of the new gene annotation we aligned and assigned RNAseq libraries from the *Nematostella* developmental time series (0-19h)[9] and compared it to the previous transcriptome. Overall we see a decrease of 2% in the fraction of unmapped reads, an increase of 10% in the fraction of reads unambiguously assigned to a gene model, a drop of 10% in the fraction of reads not assigned to any gene model and a drop of 5% in the fraction of multi mapping reads which cannot be reliably assigned to a specific gene model (EDF4b). Additionally, we aligned 27 public single cell libraries from Nematostella[10,11] and compared the results between the NVE models and the NV2 models. On average there was a 5% increase in the reads aligned to the NV2, a 10% drop in the fraction of reads aligned multiple times and a slight increase in the average number of reads per gene (EDF4c). Compared to the NVE models, the NV2 have longer UTRs which are benefitial to the mapping of the single cell transcriptome libraries from 10X Genomics platforms.

#### 2. Chromosomal repeat dynamics in the Edwardsiidae

Remarkably, the longest two pseudo-chromosomes in *Scolanthus* correspond to the 7th shortest and the shortest pseudo-chromosome in *Nematostella*, respectively. Both pseudo-chromosomes were rich in repetitive sequences in both species (Figure 3d-3). Based on this observation, we wondered whether any repeat classes were enriched in these chromosomes. We calculated z-scores based on the individual repeat families' fractional enrichment per chromosome relative to the distribution of chromosomal repeat enrichment. We found that in particular the LTR/Pao repeat class, a pan-metazoan repeat class abundant in *Drosophila* genomes, but absent from mammalian genomes[12], was enriched in both pseudo-chromosomes relative to others (z-scores 1.3 and 1.4, respectively, EDT 2-5) as well as their counterparts in *Nematostella* (z-scores 1.8 and 1.5).

#### 3. Evolution the homeobox gene clusters in early branching metazoans

The chromosome-level assembly of the *Nematostella* genome allows us not only to follow the evolution of gene linkages by comparing macrosyntenies at the genome-wide scale, but also to re-address the evolution of specific gene clusters. Some of the best-known examples of clusters of regulatory genes in Bilateria include the homeobox genes of the *Antennapedia* class (ANTP), the paired class (PRD), the TALE-class (*Irx* genes) and the SINE-class (*SIX* genes). For the ANTP genes, a single ancestral Megacluster containing the ProtoHox cluster linked to *Evx*, *Mox* and *Dlx*, the *NK/NK-like* homeobox genes, as well as the EHGbox genes (*Engrailed*, *HB9/Mnx*, *Gbx*), has been hypothesized. Upon duplication of the ProtoHox cluster, the Hox and the ParaHox clusters arose, and the Megacluster broke up into several fragments[13]. The later continuation of the work of Pollard and Holland added several genes to the “extended Hox” or “SuperHox” cluster of the Urbilaterian, which was postulated to have contained the *Hox*, *Evx*, *Dlx*, *Nedx*, *Engrailed*, *Mnx*, *Rough*, *Hex*, *Mox*[14] and *Gbx*[15]. Comparison of the genomic linkages in the early-branching chordate *Branchiostoma floridae* (Deuterostomia) and the annelid *Platynereis dumerilii* (Protostomia) then allowed to postulate that the last common ancestor of the protostomes and deuterostomes had the following four clusters of ANTP genes: the SuperHox cluster, the ParaHox cluster, the NK/NK-like cluster, and the NK2 family cluster[14]. In contrast to the earlier branching sponges, which only contain the NK/NK-like cluster of the ANTP class genes[16], *Nematostella vectensis* has the four clusters similar to that postulated for the hypothetical urbilaterian by Hui et al[17].

The *Nematostella* extended Hox cluster is located on the chromosome 2 (Figure 3; EDF 8) and consists of the immediately linked three Hox genes (*HoxC*/*Anthox7*,

*HoxDa/Anthox8a* and *HoxDb/Anthox8b,* the latter two likely being alternative splice variants of the single *HoxD/Anthox8*), *Evx*, *HoxA* (*Anthox6*) followed by eight bystander genes, and then by *Mnx* and *Rough*. Two more Hox genes, (*HoxE*/*Anthox1a* and *HoxR*/*Anthox9*) are located ~2.5Mb downstream of *Rough*, and one further Hox gene, *HoxB*/*Anthox6a*, is located ~5Mb upstream of *HoxC*. The last remaining *Nematostella* Hox gene, *HoxF*/*Anthox1*, a paralog of *HoxE/Anthox1a*, is found on chromosome 5. *Gbx* and a cluster of four *Mox* paralogs can also be found on the chromosome 2: *Gbx* is located ~0.4Mb downstream of *HoxB*, and *Mox* genes are positioned at the end of the same chromosome some 5.6Mb downstream of *HoxE*. A likely *Engrailed* ortholog *Engrailed-like* and *Dlx* are linked, but located on a different chromosome (Figure 3, EDF 8), and *Nedx* is not present in the *Nematostella* genome. This unusual "interrupted" structure of the SuperHox cluster with *Evx* in the middle and parts of the cluster far away from the *Hox*-*Evx*-*Mnx*-*Rough* core indicate some secondary rearrangement (Figure 3, EDF8). The existence of clear Hox and ParaHox genes in Cnidaria and in Bilateria suggests that the duplication of the ProtoHox cluster took place prior to the cnidarian-bilaterian split, although it is not clear, whether this Protohox cluster consisted of two or three genes[18]. Phylogenetic analyses[18–21] show very low support for the orthology of the cnidarian Hox genes with distinct groups of bilaterian Hox paralogs, which resulted in an attempt to use non-tree-based methods to establish the phylogenetic relationships between them[22]. Given that the *Nematostella* ParaHox cluster (*N. vectensis* chromosome 10; Figure 3) consists of two linked genes – an "anterior" *Gsx* and a mixed identity "non-anterior" *Xlox*/*Cdx*[18], it seems likely that Hox genes in Cnidaria and Bilateria are a result of the independent diversification of the descendants of a single "anterior" and a single "non-anterior" Hox gene of the cnidarian-bilaterian ancestor (Figure 3a), although the presence of the complete cnidarian ParaHox cluster containing Gsx, Xlox and Cdx cannot be excluded based on the evidence from a scyphozoan *Rhopilema*[23]. The putative ancestral two-gene Hox cluster was linked to *Eve*, *Mnx*, *Rough* and, possibly, *Mox*. Since the SuperHox cluster of the scyphozoan jellyfish also appears to be at least partially atomized[23], it remains unclear whether the remaining members of the bilaterian SuperHox cluster (*Gbx*, *Dlx*, *Engrailed-like*, if the latter is a true *Engrailed* ortholog) were also linked to it or got “trapped” by it in the bilaterian lineage.

Similar to the situation in Bilateria, the *Nematostella NK/NK-like* homeobox genes are concentrated in two locations: the five *NK2.2* genes form a cluster on the chromosome 2, and there is an NK cluster on the chromosome 5 (Figure 3, EDF 8-9). Within a space of less than 1Mb, there is a group of *NK6*, *NK7*, *NK3*, *NK4*, *Msx*, *NK5* and *NK1*. These are seven out of nine members of the ProtoNK cluster postulated for the protostome-deuterostome ancestor[24]. The missing two genes, *Ladybird* (*Lbx*) and *Tlx-like* (its orthology with bilaterian Tlx is not fully certain - it is suggested by BLAST but not supported by the phylogenetic analyses (EDF 8-9)), are found on the same chromosome respectively 8.2Mb and 9.3Mb upstream of *NK6*. Since in the sponge *Amphimedon* a suggested *Tlx* ortholog (although we do not find statistical support for sponge Tlx orthology in our analyses (EDF 8)) is part of the NK cluster[16], it is possible that *Tlx-like* outside of the NK cluster in *Nematostella* represents a derived rather than an ancestral trait. Alternatively, it may be that true *Tlx* genes first evolved in Bilateria. More orthologs of the bilaterian NK-like genes and possible members of the ancestral eumetazoan NK cluster[15–17,24] are found on the same chromosome at varying distances to the cluster: *Emx* (5 *Emx* genes in different locations on the chromosome), *Hlx*, *Not*, and, intriguingly, *Hex*. Multiple additional NK-class genes are located on different chromosomes, including one *Msx* copy approximately 0.34Mb upstream of the *HoxE*/*Anthox1a*.

*Hex* was proposed to be a member of the SuperHox rather than the NK cluster of the bilaterian ancestor[14]. At the same time, in the sea anemone, *Hex* is found on chromosome 5 carrying the NK cluster as well as many other NK-like genes and, notably, *HoxF/Anthox1*. The presence of the *HoxF* gene, sometimes in several copies, on the chromosome carrying the NK cluster appears to be a conserved cnidarian feature, since this is also the case in another edwardsiid sea anemone *Scolanthus*, the octocoral *Xenia* and the scyphozoan *Rhopilema*. *Hex* appears to be the single potential SuperHox cluster gene present in the ctenophore and sponge genomes[25–29]. Like the sea anemone *Hex*, the proposed *Hex* orthologue in the sponge *Amphimedon queenslandica* is part of the NK cluster[16] (although, again, we do not find strong statistical support for sponge Hex orthology in our analyses (EDF 9)). Thus, the position of *Hex* on the “NK-cluster chromosome” in *Nematostella* may represent the ancestral condition. It is also possible that *Hex* linked to the NK cluster may be a remnant of the SuperHox-NK Megacluster, which independently split into SuperHox and NK clusters in Cnidaria and Bilateria. In this scenario, cnidarian SuperHox cluster must have translocated onto what is now *Nematostella* chromosome 2 leaving *HoxF*/*Anthox1* and *Hex* behind; and bilaterian SuperHox cluster must have translocated into its current position taking all the SuperHox cluster genes (including *Hex*) with it. However, it is also possible that *Hex* became translocated into the SuperHox cluster from the NK cluster early in the bilaterian evolution, and the placement of the sea anemone *HoxF*/*Anthox1* on the chromosome carrying the NK cluster does not represent its ancestral position but is a result of the duplication of the “non-anterior” Hox gene and its translocation out of the SuperHox cluster at some point during the evolution of Cnidaria. This second option seems to be a much better fit to the results of our comparison of the chromosomal arrangement of the extended Hox cluster and the NK-like genes, as well as with the macrosynteny analysis. The latter shows that syntenic blocks from the SuperHox-containing *Nematostella* chromosome 2 can be found on the *Branchiostoma* chromosomes 1, 10 and 17 (where the amphioxus SuperHox cluster resides). In contrast, syntenic blocks from the NK cluster containing *Nematostella* chromosome 5 are all found on the *Branchiostoma* chromosome 3 (where the disarranged amphioxus *NK* genes are located). Moreover, clear *Hex* orthologs are members of the NK cluster not only in the sea anemone, but also in deuterostome bilaterians such as the cephalochordate *Branchiostoma* and the ambulacrarian *Saccoglossus*[30]. Thus, currently we do not find support for the SuperHox-NK Megacluster hypothesis. Taken together, we propose that the last common ancestor of Cnidaria and Bilateria possessed an NK-cluster on a chromosome different from the one carrying the SuperHox cluster, and a separate NK2.2-like cluster, which might have been on the same chromosome as the SuperHox cluster (Figure 3a). The later notion is supported by the localization of *Branchiostoma NK2.1* and *NK2.2* on the same chromosome as the Six gene cluster, *Goosecoid* and *Otx* in *Branchiostoma* – similar to the situation in *Nematostella* (Figure 3e and see also text below).

In addition to the previously established clustering of the PRD-class *Homeobrain-Rx-Orthopedia* conserved between cnidarians, bilaterians and placozoans[30,31], we find several other more distant linkages, which might be nonetheless meaningful. For example, similar to the deuterostome situation[30,32], LIM class gene *Isl* is linked to the *Rx-Orthopedia*, however, at a distance of approximately 4.7Mb. SINE-class genes *Six3/6*, *Six1/2* and *Six4/5* are clustered in deuterostomes either closely like in hemichordates[30] and cephalochordates or with several Mb in between, like in vertebrates. The order of the genes in this cluster appears to be highly conserved: Six3/6 > Six1/2 > Six4/5. The same order of the *Six* genes we find on the *Nematostella* chromosome 2. Another notable cluster is the so-called "pharyngeal cluster", which was considered deuterostome-specific[30]. In deuterostomes, it is composed of the *NK2.1* and *NK2.2* genes closely linked with the non-homeobox transcription factor *FoxA* and *Pax1/9* and two non-transcription factor genes *Mipol1* and *Slc25A21*. In *Nematostella*, the five *NK2.2* genes, *Mipol1* and *FoxA* are all located on chromosome 2. *Mipol1* is located between the immediately linked *NK2.2E*, *NK2.2D*, *NK2.2C* and *NK2.2A*, and the more distant *NK2.2B*, while *FoxA* is approximately 1.6Mb further downstream. We found no orthologs of *Pax1/9* and *Slc25A21* in *Nematostella*. Notably, similar to deuterostomes, among the five *Nematostella NK2.2* genes, only *NK2.2A* is not expressed in the pharynx[33], and *FoxA* is one of the most conserved pharyngeal markers in *Nematostella*[34,35]. The location of a selection of other homeobox proteins including the aforementioned PRD, TALE and LIM classes is provided in the EDT 6.

#### 4. Ultraconserved Non-coding Elements

We were interested to find out how many, and to what extent, Ultraconserved Non-Coding Elements (UCNEs) are present in Edwardsiidae. We adopted criteria previously used to detect UCNEs between chicken and humans[36] and found 145 regions in the *Nematostella* genome that were highly conserved with *Scolanthus* (EDT 7). One hundred sixteen (116) of these regions fell into 37 syntenic clusters of at most 500 kb intervening gaps, and the remaining 29 were singletons. Several such clusters were close to NK paralogs, such as one containing 12 UCNEs and spanning 70 kb surrounding the NK3-4 cluster on chromosome 5 and three UCNEs upstream of the NK2.2 cluster on chromosome 2, a pattern previously reported in vertebrates[32]. Additionally, we detected a single UCNE neighboring the *PaxC* gene. While Pax-associated UCNEs have been previously reported in vertebrates[37–39], this neural developmental gene appears to have arisen from a cnidarian-specific duplication[40], implying that the accompanying UCNE also arose independently. On the other hand, no UCNEs are found near the edwardsiid *Irx* gene, despite their ubiquity among Bilateria[41–43]. Likewise, our stringent criteria did not detect the previously reported UCNE at the 3' end of the SoxB2 gene, as previously reported[44], suggesting that more relaxed parameters might reveal additional conserved elements shared between more distantly related species.

#### 5. SIMRBase Genome Browser

We are providing a comprehensive, maintained and community-oriented genomic resource for the *N. vectensis* genome. Genome browsers, gene pages, gene searches, and BLAST interfaces for each species are available at https://simrbase.stowers.org. Users may query genes by  name, ID, and genome location and gene ontology (GO) to view data aligned to the genomes, such as transcriptomes, RNAseq and protein homologs in the genome browsers. Outside the browsers, more detailed searches can be performed using keywords from a collection of precomputed sequence similarity searches and protein domain predictions.

#### 6. Inference of ancestral linkage groups

We compared the chromosomal content and order of *Nematostella vectensis* to metazoan genomes including Porifera, other Cnidaria, Ecdysozoa, Spiralia and Chordata. We were able to observe differential rearrangement and conservation of genes among chromosomal pairs, and we hypothesized that some of these chromosomes shared among *Nematostella* and many other genomes in fact derived from the same ancestral linkage group. In order to directly determine the extent to which these ancestral linkage groups are shared, we took a graph-based approach for the inference of both orthologous groups and macrosyntenic links.

In order to determine orthology groups, we used the OMA standalone algorithm[45] to determine cliques of orthology within our large set of genomes. We chose this in order to avoid the misidentification of genes which are either paralogous and would not be expected to have retained their chromosomal linkage or are partial matches to loci originating from duplicated regions arising from misassemblies. This resulted in several large orthology groups, summarized in EDF 13.

The orthology groups from the previous step were used to construct a macrosynteny graph whose nodes were the orthology groups. We added edges to the graph for each pair of groups that occur on the same chromosome or scaffold in two different species. We then endeavored to determine ancestral linkage groups (ALGs) among critical internal nodes of the metazoan tree and the root, using subgraphs of the macrosynteny graph of species contained in each clade. Specifically, we used the genomes of *N. vectensis, E. pallida, A. millepora, Xenia sp., A. aurita, R. esculentum, C. hemisphaerica* and *H. vulgaris* to determine the cnidarian ancestral linkage groups, the genomes of *C. rotundicauda, T. corallinus, D. melanogaster, P. yessoensis* and *L. oculatus* to determine the bilaterian ancestral linkage groups and both of the above groups combined with *T. adhaerens, E. muelleri* and *A. queenslandica* to determine the metazoan ancestral linkage groups.

Our approach to determining the groups from the graph is centered around the Leiden algorithm[46] as implemented in igraph version 1.2.5[47], which finds well-connected communities. The output of the algorithm is a “partition” of the input vertices into a number of groups. In our case, this translates to groups of genes which are frequently detected on the same chromosomes in corresponding genomes. We based our approach around the modularity optimization function, which maximizes the number of edges within communities as compared to the expected number of edges[48]. In the case of the Leiden algorithm, a “resolution” parameter defines the granularity of community detection, in other words, the number of communities detected.

As the number of ancestral linkage groups is critical to the interpretation of the result, we chose to optimize the number of clusters detected based on the reported modularity of various values of the resolution parameter $\gamma$. Instead of fixing the value, we initially ran the Leiden algorithm 100 times for each $\gamma$ value between 0.5-1.5, by steps of 0.05. The ranks were then used to generate a stepwise function $f(\gamma) = \frac{\Sigma_{\gamma_i=\gamma} r_i}{N}$ where $r_i$ is the rank of partition $i$ with respect to modularity and $\gamma_i$ is the gamma value used for partition $i$, and $N$ is the total number of partitions. This was transformed into a smooth local regression function generated using the locfit algorithm[49] (EDF 14).

Optimizing modularity is an NP-hard problem[50] (i.e., there is no known solution whose time scales polynomially with the number of nodes in the graph), and therefore the Leiden algorithm is an approximation to the optimal solution. In order to find high-confidence communities, we chose to use a consensus approach. We sampled the above-determined distribution of $\gamma$ parameters 1000 times to generate a pool of partitions by running the Leiden algorithm with the sampled $\gamma$ values. The pool of 1000 partitions was then used to determine the consensus among the many runs of the Leiden algorithm. The consensus number of clusters $k$ was determined as the maximum of $f(k) = \frac{\Sigma_{k_i=k} m_i}{N_k}$ where $m_i$ is the modularity of partition $i$ and $N_k$ is the number of partitions with $k$ groups. We synthesized the results of all 1000 partitions to a consensus using the algorithm implied by the first model of Gordon and Vichi[51].
